# A unified mechanism for spatiotemporal patterns in somitogenesis

**DOI:** 10.1101/2020.03.28.013284

**Authors:** Chandrashekar Kuyyamudi, Shakti N. Menon, Sitabhra Sinha

## Abstract

Somitogenesis, the process of body segmentation during embryonic development, exhibits a key set of features that is conserved across all vertebrate species despite differences in the detailed mechanisms. Prior to the formation of somites along the pre-somitic mesoderm (PSM), periodic expression of clock genes is observed in its constituent cells. As the PSM expands through the addition of new cells at its posterior, the oscillations in the cells closer to the anterior cease and eventually lead to formation of rostral and caudal halves of the somites. This pattern formation is believed to be coordinated by interactions between neighboring cells via receptor-ligand coupling. However, the mechanism underlying the transition from synchronized oscillations to traveling waves and subsequent arrest of activity, followed by the appearance of polarized somites, has not yet been established. In this paper we have proposed a unified mechanism that reproduces the sequence of dynamical transitions observed during somitogenesis by combining the local interactions mediated via Notch-Delta intercellular coupling with global spatial heterogeneity introduced through a morphogen gradient that is known to occur along the anteroposterior axis of the growing PSM. Our model provides a framework that integrates a boundary-organized pattern formation mechanism, which uses positional information provided by a morphogen gradient, with the coupling-mediated self-organized emergence of collective dynamics, to explain the processes that lead to segmentation.

## I. INTRODUCTION

Somitogenesis is the process of formation of somites, which are the modular building blocks of all vertebrate bodies [1–3]. Somites compose bilaterally symmetric segments that are formed in the paraxial, or pre-somitic, mesoderm (PSM) of developing embryos as the body axis itself elongates [4]. Analogous processes have been implicated in the body segmentation of some invertebrates [5, 6]. Although there is great variability across species in terms of the number of somites, the mean size of a somite and the duration over which they are formed, nonetheless a conserved set of features characterizing somitogenesis is seen across these species [7]. A general conceptual model for explaining these core features is provided by the Clock and Wavefront (CW) framework proposed by Cooke and Zeeman in 1976 [8], wherein the PSM was assumed to comprise cellular oscillators (clocks) which are each arrested at their instantaneous state of activity upon encountering a wavefront that moves from the anterior to posterior of the PSM [9–16]. This provides a route for translating a sequence of temporal activity into spatial patterns [17]. In order to construct an explicit mechanism embodying the CW framework, we need to disaggregate its components that operate at different scales, namely, (i) the cellular scale at which oscillations occur, (ii) the inter-cellular scale at which contact-mediated signaling takes place, and (iii) the scale of the PSM across which morphogen gradients form and act as the environment that could modulate the intercellular interactions. This resonates with the proposal of Oates [18] to view the CW framework as a three-tier process. In the bottom tier, we observe oscillations at the level of a single cell in the PSM, arising from the periodic expression of clock genes [19–26]. The middle tier describes the mechanism by which the cellular oscillators coordinate their activity with that of their neighbors. This occurs through juxtacrine signaling brought about by interactions between Notch receptors and Delta ligands [22, 27–38]. Indeed, several earlier models have explored the role of Notch-Delta coupling in bringing about robust synchronization between the oscillators [39–43]. Finally, processes that bring about the slowing down (and eventual termination) of the oscillations [19, 32, 44], and the subsequent differentiation of the cells into rostral and caudal halves of the somites, constitute the top tier. The model proposed here integrates these different scales by investigating genetic oscillators interacting via Notch-Delta coupling whose strength is modulated in a position-dependent manner due to a morphogen concentration gradient along the anteroposterior (AP) axis of the PSM. This provides an unified framework for explaining the dynamical transitions observed during somitogenesis.

As the PSM expands along the AP axis through the addition of new cells at the tail [4], it is reasonable to restrict our attention to the process of somitogenesis taking place along a one-dimensional array of coupled cells in the PSM. From the perspective of our modeling which explicitly investigates the role of morphogen gradient in coordinating somite formation, the array of cells is considered to be aligned along the AP axis, as the various morphogens that are expressed along the PSM are known to form concentration gradients along this axis [45]. These primarily include molecules belonging to the FGF [46, 47], RA [48, 49] and Wnt families [50–52]. Even though the role of gradients on the overall dynamics has been explored [53–56], there is to date no consensus as to the explicit mechanism through which they contribute to somite formation. Our model demonstrates that if the morphogen gradient is considered to regulate the strength with which adjacent cells interact, it can lead to qualitatively different kinds of dynamics along the PSM in a threshold-dependent manner [Fig. 1 (a)]. Unlike the conventional boundary-organized pattern formation paradigm [57], here the morphogen gradient does not determine the cell fate so much as affect the local dynamics such that, at any point in time, there can be qualitatively different types of dynamics occurring in separate locations in the PSM. Thus, our results help address an open question as to how morphogen gradients influence the collective dynamics of the cellular oscillators, leading to their eventual fates as rostral and caudal halves of the mature polarized somites.

**FIG. 1.**
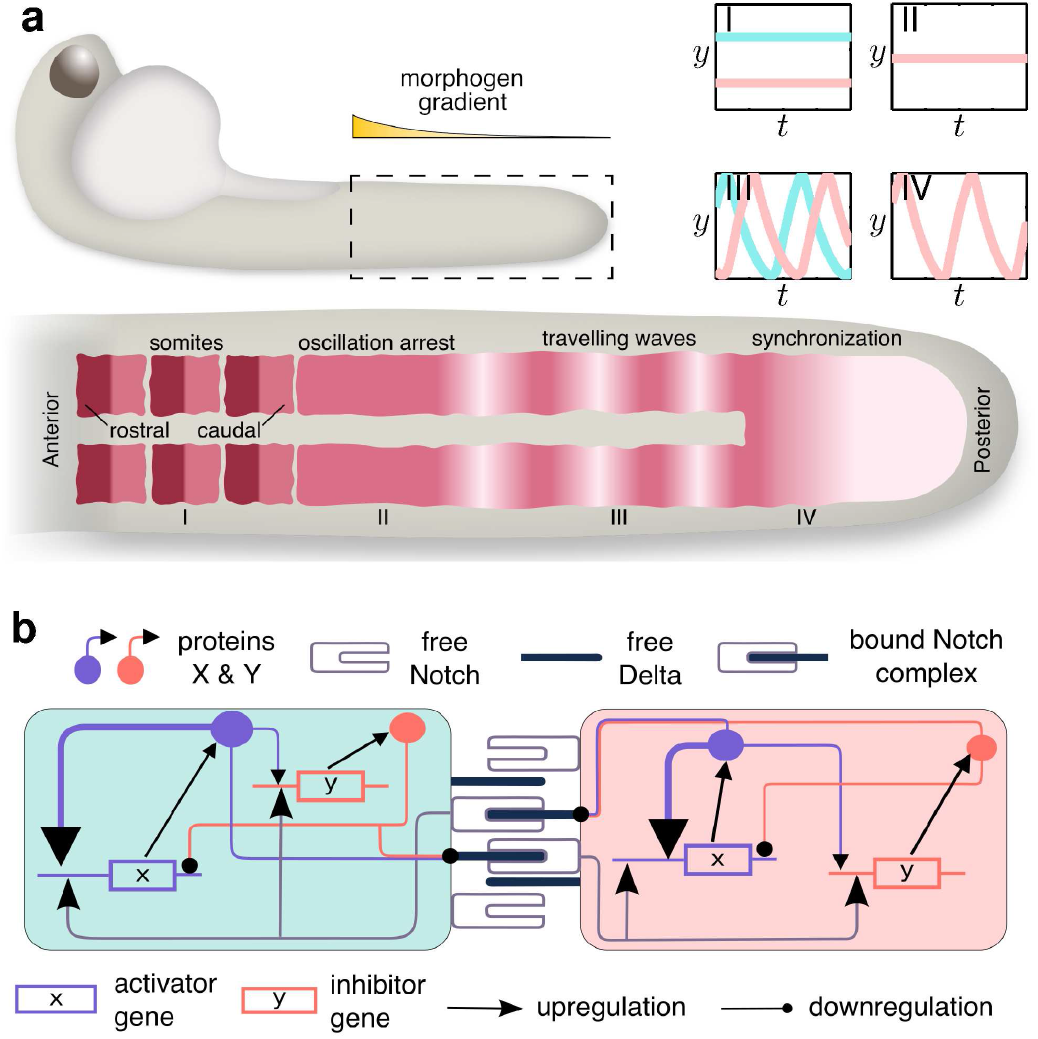
The key dynamical features of somitogenesis (top) that are reproduced in our mathematical model (bottom). (a) Schematic diagram depicting the zebrafish pre-somitic mesoderm (PSM) and the dynamical states exhibited by the cells over the course of somitogenesis. The box outlined with broken lines represents the posterior region which exhibits a spatial gradient in morphogen concentration, represented by the exponentially decaying profile shown above the box. The dynamics of the cells at the tail (region IV) are characterized by exact synchronization of the periodic variation in gene expression. As the cells move towards the anterior, they first exhibit travelling waves of gene expression (III), followed by arrest of the oscillations (II). Eventually the cells differentiate (I) into alternating bands corresponding to rostral and caudal halves of the mature somites. The figures in the insets at the right display typical time series in each of the regimes I-IV of the inhibitor gene expression for two neighboring cells coupled to each other. (b) Schematic diagram describing the interaction between two cells via Notch-Delta coupling. In general, each cell has Notch receptors, as well as, Delta ligands that bind to them. The *cis* form of the binding leads to the loss of receptors and ligands without resulting in any downstream signaling, whereas *trans* Notch-Delta binding gives rise to cleavage of the Notch Intra-Cellular Domain (NICD). The latter acts as a transcription factor (TF) for the downstream activator (*x*) and inhibitor (*y*) genes which are the essential constituents of each cellular oscillator.

## II. METHODS

### A. Modeling genetic oscillators interacting via Notch-Delta coupling

Several experiments have established that the cells in the presomitic mesoderm (PSM) have “clock” genes whose expression levels oscillate [19–24, 26]. In our model, we consider a generic two component genetic oscillator comprising an activator gene (*x*), which upregulates its own expression, as well as that of an inhibitor gene (*y*), which suppresses the expression of the activator gene [58]. The dynamics of this two-component oscillator can be expressed in terms of a pair of rate equations describing the change in concentrations of the protein products *X* and *Y* of genes *x* and *y*, respectively. The model parameters are chosen such that *X* and *Y* exhibit limit cycle oscillations.

As the communication between cells in the PSM is crucial in mediating their collective behavior during somitogenesis, we couple the dynamics of the clock genes of neighboring cells. Experiments have established the role of the Notch-Delta juxtacrine signaling pathway in mediating the interaction between cells that are in physical contact with each other [22, 27–36]. In general, each cell has both Notch receptors as well as Delta ligands on their surface. A Notch receptor on cell *i* which is bound to a Delta ligand belonging to a neighboring cell j (i.e., *trans* binding) leads to the cleavage of the Notch intracellular domain (NICD) that will act as transcription factor for downstream genes in cell *i* [59, 60]. In our model we assume that the NICD upregulates the expression of both the clock genes, while the gene products *X, Y* suppress the production of Delta ligands by the cell [39, 41]. We describe the Notch-Delta signaling mechanism through the coupled dynamics of (i) the free (unbound) Notch receptor concentration (*N*), (ii) the free Delta ligand (*D*) and (iii) the NICD which is released as a result of *trans* binding (*N^b^*). Thus, the dynamics of a cell *i* coupled to its neighbors through Notch-Delta signaling is described by the following set of equations:

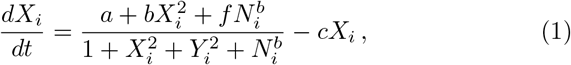

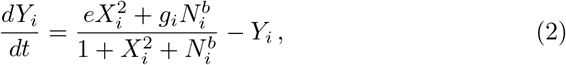

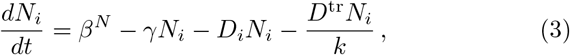

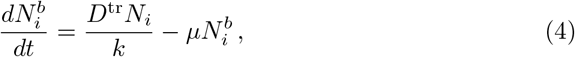

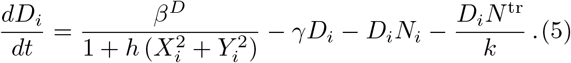

Here the terms *D^tr^* and *N^tr^* are the mean values of *D_j_* and *N_j_* over all neighboring cells *j* to which *i* is coupled through *trans*-binding. While the values of the model parameters can, in general, vary across cells, we restrict our attention to the variation of the coupling parameter *g* (subscripted with the cell index in Eqns. (1-5)), which determines the strength of upregulation of *y* by the NICD (*N^b^*). This allows us to investigate the role of spatial heterogeneity imposed by the gradient of morphogen concentration along the anteroposterior (AP) axis of the PSM.

### B. Dynamical evolution of the morphogen gradient

Our model focuses on the behavior of a contiguous segment of cells of length *ℓ* in the PSM with a morphogen source located at its anterior. If the strength of the source is constant in time, it would have resulted in the gradient becoming progressively less steep as the PSM expands. This dilution is countered by the net increase in the strength of the source through the secretion of morphogen by the newly matured somites. Thus, as the PSM expands due to addition of cells at the posterior tail, we can view the segment under consideration as effectively flowing up a morphogen gradient along the AP axis [18]. We choose a segment of *N* cells with a spatial extent *ℓ* that is initially located (*t* = 0) at the posterior end of the PSM, i.e., at the lower end of the gradient, and follow its evolution upto a time period *T*. As mentioned above, the effect of the varying morphogen concentration on the dynamics of the cells is introduced via the coupling parameter *g*. Specifically, we assume that the value of *g* at each site is proportional to the corresponding morphogen concentration, yielding an exponentially decaying gradient of *g* across the AP axis: *g_i_*(*t*) = *g_min_* exp (*λ_g_x_i_*(*t*)). The steepness of the gradient is quantified by *λ_g_*, which is a function of *T*, as well as *g*_max_ and *g*_min_, which are the values of *g* at the anterior and posterior ends of the PSM, respectively, viz., *λ_g_* = ln(*g*_max_/*g*_min_)/*T*. We assume that the effective flow of the segment of cells along the AP axis occurs at an uniform rate. This can be taken to be unity without loss of generality by appropriate choice of time unit. Thus, the instantaneous position *x_i_*(*t*) along the gradient of the ith cell in the segment is given by *x_i_*(*t*) = *t* + (*ℓ*/*N*)(*i* − 1), with the initial condition as *x_i_*(*t* = 0) = 0.

## III. RESULTS

In our simulations, we have considered the PSM to comprise cells, each of which exhibits oscillating gene expression. We assume a minimal model for the genetic oscillator consisting of two clock genes, one activatory and the other inhibitory, whose products correspond to fate determining proteins (see Methods). The oscillations of neighboring cells influence each other through Notch-Delta inter-cellular coupling [Fig. 1 (b)]. We have explicitly verified that incorporating delay in the contact mediated signaling does not alter our results qualitatively (see SI). Considering the simplest setting, viz., a pair of adjacent cells, which allows us to investigate the effect of coupling on the collective dynamics, we see from Fig. 2 that the system can exhibit a wide range of spatio-temporal patterns. These can be classified systematically through the use of quantitative measures (see Supplementary Information for details). We focus on the patterns that can be immediately interpreted in the context of somitogenesis: (i) Inhomogeneous Steady States (ISS), (ii) Homogeneous Steady States (HSS), (iii) Anti-Phase Synchronization (APS) and (iv) Exact Synchronization (ES) [shown schematically as insets of Fig. 1 (a)]. The range of values of the coupling-related parameters *f*, *g* and *h* over which these patterns are observed in a pair of coupled cells are shown in Fig. 2. The parameters f and *g* govern the strength with which the Notch intra-cellular domain (*N^b^*) regulates the activatory and inhibitory clock genes, respectively, while *h* is related to the intensity of repression of the Delta ligand (*D*) by each of the clock genes. As inter-cellular coupling is believed to be responsible for the synchronized activity of cells in the initial stage of somitogenesis [61], we note that the dynamical regime corresponding to ES occurs for low *g* and intermediate values of *f*, with the region increasing in size for larger *h*.

**FIG. 2.**
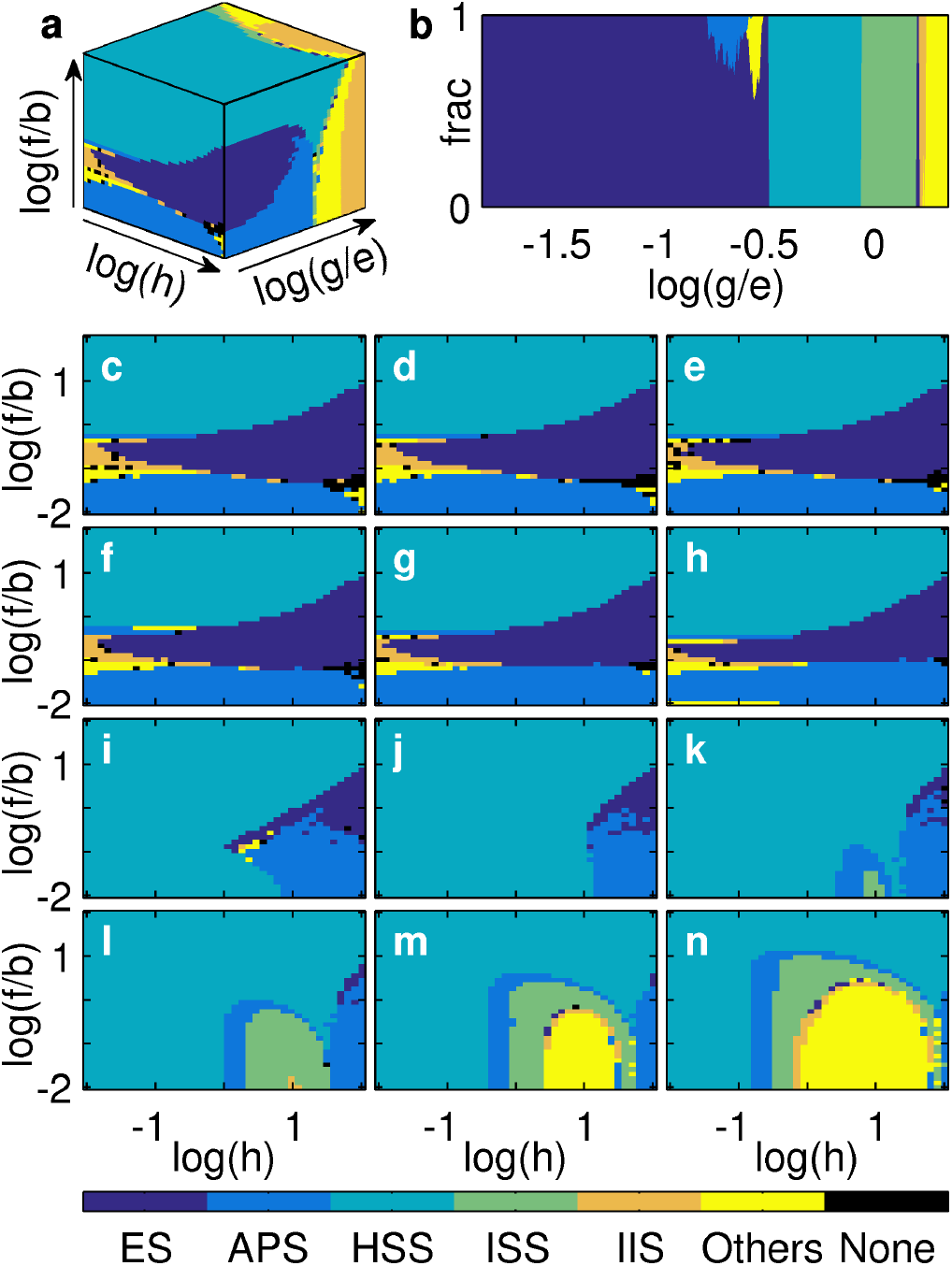
Transitions between patterns representing different stages in somitogenesis seen in the collective dynamics of a pair of cells on varying the parameters governing the strength of Notch-Delta coupling in our model. (a) Schematic diagram of the three-dimensional space spanned by the coupling parameters (*f,g,h*), scaled by the relevant kinetic parameters of the individual oscillators. Note the logarithmic scale used for the ranges of the parameters. (b) The variation with *g* of the relative frequency of occurrence of patterns belonging to each of six distinct categories, viz., Exact Synchronization (ES), Anti-Phase Synchronization (APS), Homogeneous Steady States (HSS), Inhomogeneous Steady States (ISS), Inhomogeneous In-phase Synchronization (IIS) and Others. The values of the parameters *f/b* and *h* are fixed at 0.25 and 4, respectively. (c-n) The most commonly occurring dynamical patterns (i.e., obtained for > 50% of all initial conditions used) that are seen for different values of *f/b* and *h* (varying over four order of magnitude) for 12 equally spaced values of log(*g/e*) between −1.89 (panel c) and 0.36 (panel n). While HSS is the most common pattern seen over this range of parameter values, focusing on how the occurrence frequency of ES varies with *f*, *g*, *h* indicates where a transition from synchronization to time-invariant behavior may be achieved.

Spatial variation in these coupling parameters across the PSM can arise through heterogeneity in the underlying morphogen concentrations. It is known that the morphogens RA, Wnt and FGF are differentially expressed along the PSM, exhibiting monotonically varying concentration gradients having peaks at the posterior (for FGF and Wnt) or anterior (for RA) ends [51, 62, 63]. Experiments on several vertebrate species have shown that high concentrations of RA initiate differentiation, while increased levels of Wnt and FGF, which are known to play a role in sustaining oscillations in the expression of the clock genes, impede the formation of mature somites [46, 49, 64, 65]. Thus, as our goal is to explicate the mechanisms driving the differentiation of individual cells into anterior and posterior halves of the mature somites, we focus on the role of RA on the collective dynamics of cells in the PSM. We incorporate the effect of this morphogen in the spatial variation of the coupling parameter *g* which is assumed to exponentially decay from the anterior to the posterior end of the domain. Such a profile will naturally arise if the morphogen diffuses from a source located at the anterior and is degraded at a constant rate across space [66–68]. We note that qualitatively similar results are obtained if gradients are introduced in either *f* or *h* instead of *g*. The effect of Wnt and FGF are implicitly accounted for by the endo-geneously oscillating system of two coupled clock genes that we consider in our model.

Introducing heterogeneity through the coupling parameter in the pair of adjacent cells considered earlier, we observe that qualitatively similar spatio-temporal patterns to those observed in Fig. 2 are obtained (see Supplementary Information, Fig. S3). On varying the mean value of the coupling 〈*g*〉 = (*g*_1_ + *g*_2_)/2, which effectively represents the location of these cells on the PSM, and the steepness of the gradient |*g*_1_ − *g*_2_| [Fig. 3 (a)], the range of 〈*g*〉 over which ES is seen (corresponding to the region proximal to the posterior end of the PSM) does not appear to change appreciably on increasing |*g*_1_ − *g*_2_| [Fig. 3 (b)]. Above a critical value of 〈*g*〉 which is independent of the gradient, the activities of the cells are arrested at *Y*_1_ and *Y*_2_, respectively, with the gap Δ = |*Y*_1_ − *Y*_2_| becoming larger as we move towards the anterior end, corresponding to increasing 〈*g*〉 [Fig. 3 (c)].

**FIG. 3.**
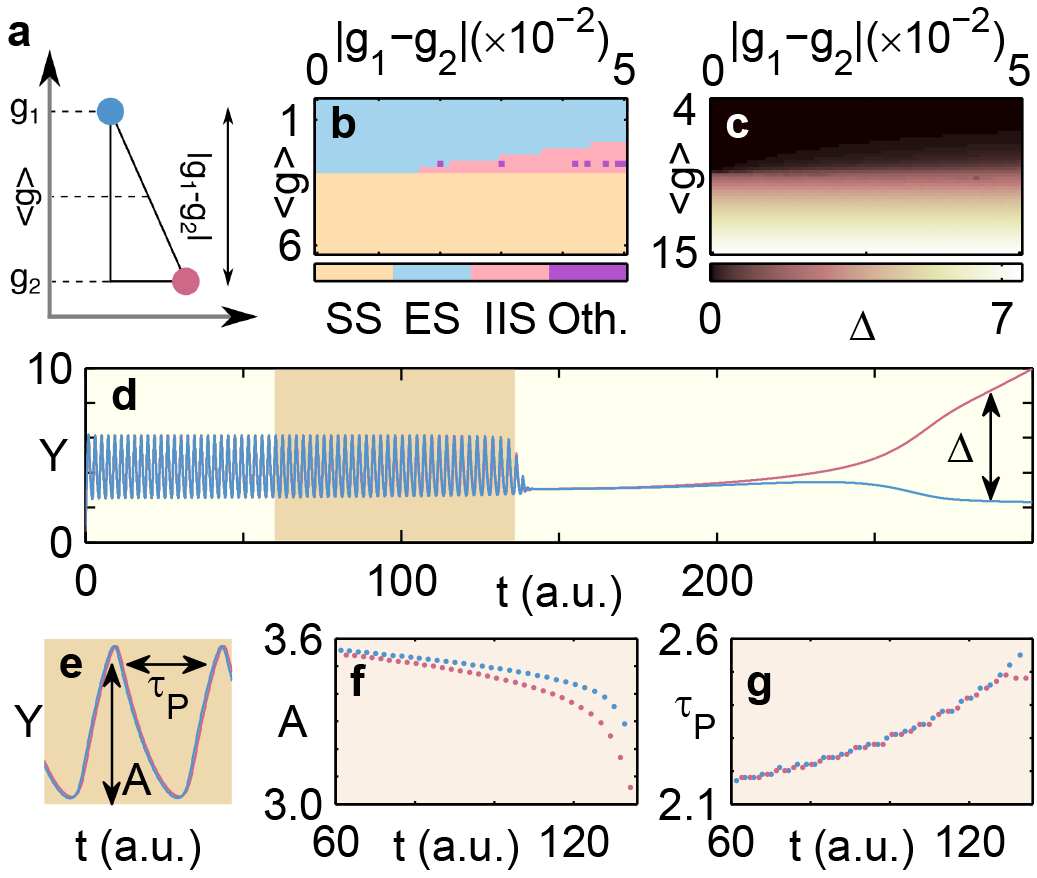
Incorporating a morphogen gradient by varying the Notch-Delta coupling parameter *g* of adjacent oscillators reproduces the temporal sequence of patterns observed during somitogenesis. (a) Schematic representation of the spatial variation of g, resulting from the different concentrations of a morphogen sensed by neighboring cells. The steepness of the gradient is quantified by |*g*_1_ − *g*_2_|, the difference in the values *g*_1,2_ for the two oscillators, while their location on the gradient is determined by the mean (*g*). (b) The most commonly occurring dynamical patterns (i.e., obtained for > 50% of all initial conditions used) on varying |*g*_1_ − *g*_2_| and 〈*g*〉. The steady state (SS) region, representing both HSS and ISS patterns, spans a large area across parameter space. In this region, the states of the adjacent cells, which are characterized by the protein concentration *Y*, converge to fixed points *Y*_1,2_. (c) The difference between the steady state concentrations of the adjacent cells, Δ = |*Y*_1_ − *Y*_2_|, increases with 〈*g*〉 as one effectively moves up the morphogen gradient, while being relatively unaffected by the steepness |*g*_1_ − *g*_2_|. (d-g) The dynamical consequences of a morphogen gradient with an exponentially decaying concentration profile. (d) As a pair of coupled cells gradually move upstream of the gradient, resulting in an increase of *g*_1,2_ over time *t*, their collective behavior (represented by *Y*) converges from initially synchronized oscillations (ES) to an inhomogeneous steady state (ISS, characterized by finite values of Δ) at long times [see SI, Fig. S5]. The changes occurring in the system during the transition from oscillations to steady state behavior (shaded region) can be quantitatively investigated by focusing on how the amplitude *A* and period *τ_p_* of the oscillations (see panel e) change over time. (f-g) As cells move upstream of the morphogen gradient (corresponding to a progression from posterior to anterior regions in the PSM), the model exhibits decreasing *A* (f) and increasing *τ_p_* (g). This is consistent with a key experimental observation, viz., shortening and slowing of oscillations as cells approach the anterior end of the PSM, during somitogenesis [18].

As explained in the Methods, over the course of development, the PSM expands through new cells being added to its posterior end, such that the existing cells progressively encounter increasing values of the morphogen concentration. Modeling this time-evolution as an effective flow of the segment of adjacent cells along the gradient in *g*, we observe that a transient phase of ES is followed by desynchronization and subsequent attenuation of the oscillations, eventually leading to a separation of the steady states of the two cells [Fig. 3 (d)]. The gap Δ between the steady states increases with time, giving rise to a pronounced ISS state. Immediately preceding the arrest of periodic activity, we observe that the amplitude *A* and period *τ_P_* [Fig. 3 (e)] of the oscillations show characteristic changes that are in agreement with experimental observations of somitogenesis. Specifically, the amplitude decreases [Fig. 3 (f)] while the period increases [Fig. 3 (g)] with time, similar to what has been reported in Refs. [18, 69].

Having seen that a pair of contiguous cells can indeed converge to markedly different steady state values of their clock gene expressions, we now investigate the generalization to a spatially extended segment of length *ℓ* in the growing PSM subject to a morphogen gradient [varying from *g*_min_ to *g*_max_ as shown schematically above Fig. 4 (a)]. As the principal variation of the morphogen concentration occurs along the AP axis of the PSM, we restrict our focus to a one-dimensional array of cells aligned along this axis. As new cells are added to the posterior of the PSM over time, the relative position of the segment of cells under consideration shifts along the morphogen gradient from the posterior to the anterior. We note that had there been a temporally invariant source of morphogen at the anterior, the expansion of the PSM would have resulted in a dilution of the gradient. However, newly matured somites at the anterior end serve as additional sources of the morphogen over time [70–72], thereby ensuring that each cell experiences an exponential increase in the morphogen concentration (see SI). This is reproduced in our model by the segment effectively flowing up the morphogen gradient (as discussed in Methods).

**FIG. 4.**
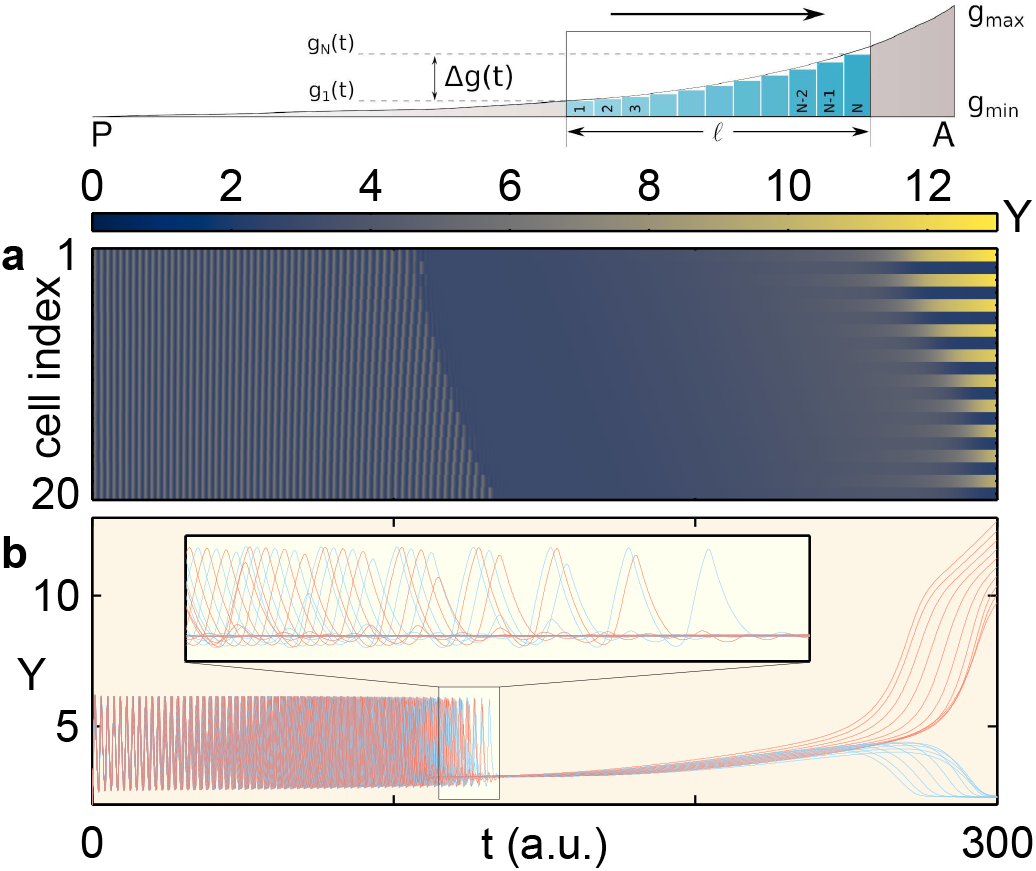
Collective dynamics of a cellular array responding to an exponential gradient of morphogen concentration reproduces the spatio-temporal evolution of PSM activity seen during somitogenesis. The transition from progenitor cells in the posterior (P) to maturity at the anterior (A) of a segment of length *ℓ* comprising *N* cells in the PSM viewed as a flow upstream (moving window in the schematic on top) over a period of time *T* along the exponential profile of the parameter *g*, decaying from gmax to *g_min_*, reflecting the morphogen gradient. Different cells in the window sense different morphogen concentrations, whose values change over time. This is incorporated in terms of the time-dependent gradient Δ*g*(*t*), with *g*_1_(*t*) and *g_N_*(*t*) being the values of the coupling parameter *g* at the anterior and posterior ends of the window at time *t*, respectively. The resulting change in the activity of a segment comprising *N* = 20 coupled cells, as it moves from P to A, is shown in (a). The system initially exhibits synchronized oscillations across the segment but, as a consequence of the gradient, a phase lag develops between adjacent cells (as seen in the time series in panel b). This results in a wave-like propagation of the peak expression from the anterior to the posterior end of the segment in each cycle. Subsequently, the oscillations reduce in intensity leading to a homogeneous steady state (HSS), a magnified view of the transition being shown in the inset of panel (b). Eventually, the system converges to an inhomogeneous steady state (ISS), with adjacent cells attaining different fates characterized by alternating high and low values of the protein concentration *Y* [cells with odd and even indices on the segment are shown using different colors in panel (b)]. Note that the system exhibits the entire range of patterns shown in the schematic in Fig. 1 (a), with the temporally invariant spatial pattern observed at the anterior end of the PSM resembling that of alternating rostral and caudal halves of somites. Results shown are for parameter values of *g*_max_ = 15.0, *g*_min_(= *g*_0_) = 1.0 and *T* = 300.0 a.u.

As shown in Fig. 4 (a), for a range of values of the parameters *g*_max_, *g*_min_ and *ℓ*, the cells display a short-lived ES pattern, which is followed by the development of a phase lag between adjacent cells [as can be seen in the inset of Fig. 4 (b)]. This is analogous to the appearance of a small phase difference between the pair of oscillators described earlier, and manifests as a travelling wave that propagates along the segment. As the cells move further up the gradient, the oscillations subside and the segment exhibits a transient homogeneous state, which is consistent with the experimental observation of cessation of expression dynamics before the mesoderm is segmented into somites [73]. This eventually gives way to a heterogeneous steady state characterized by adjacent cells having alternating high and low clock gene expressions. The gap between these high and low values increases with time to eventually produce a distinctive pattern that resemble the stripes that arise due to polarization of each somite into rostral and caudal halves [as seen for large *t* in both Figs. 4 (a) and (b)]. In this asymptotic steady state, the separation between the high and low values for clock gene expression is greater than the amplitude of the oscillations seen at lower values of *t*. Thus, we can reproduce the entire sequence of dynamical transitions observed in the PSM during somitogenesis through a model incorporating an array of oscillators that interact via Notch-Delta signaling while “moving up” a morphogen gradient.

## IV. DISCUSSION

Somitogenesis is seen across all vertebrates, and recent evidence implies that mechanisms underlying it could have analogues even in segmentation of invertebrates, such as arthropods [5, 74]. It would appear that there is an invariant set of mechanisms responsible for this process, that differ only in terms of the specific identities of the contributing molecular players across species. Thus, somitogenesis would in general involve (i) a cellular “clock”, (ii) means by which neighboring clocks communicate, and (iii) a spatial gradient of signaling molecules, which introduces heterogeneity in the interactions between the clocks. We have shown here that incorporating these three elements in a model of a PSM, that grows through the addition of cells at the posterior, reproduces the sequence of invariant dynamical transitions seen in somitogenesis. These include the phenomenon of somite polarization, provided that the genes determining the fates of cells constituting the rostral and caudal halves of the mature somites [e.g., *mesp-b* in zebrafish [75–77]] are located downstream of the clock genes.

While the roles of interacting clocks and that of morphogen gradients have been investigated individually in earlier studies, we provide here a framework to understand how these two work in tandem to give rise to the key features associated with somitogenesis. In particular, our results shed light on the significance of the steepness of the morphogen gradient. For instance, we may consider the consequences of a reduction in the steepness leading to a linear profile for the morphogen gradient which can arise, for example, when the degradation rate is negligible. On replacing the exponential morphogen gradient in our model with a linear one, we observe a very long-lived transient state before the system converges to an inhomogeneous steady state. This therefore suggests that exponential gradients allow relatively rapid switching between qualitatively distinct dynamical regimes [see SI for results with linear gradient]. Hence, by varying the steepness of the RA gradient experimentally it should be possible to determine how the time required for maturation changes as a consequence. This is especially true in the case of the time interval between cessation of oscillations and the polarization of the somites. As intercellular coupling is also known to regulate the period of the segmentation clock [78], it is possible that introducing other morphogen gradients (such as, Wnt and FGF), that influence the strength of the coupling, can explain variations in the rate at which somites form over time. Furthermore, the core assumption of our model, namely that Notch-Delta coupling plays a crucial role in regulating somitogenesis in the presence of a morphogen gradient can be probed in experimental systems where Notch signaling has been arrested.

## ACKNOWLEDGMENTS

We would like to thank Krishnan Iyer, Jose Negrete Jr, Shubha Tole and Vikas Trivedi for valuable suggestions. SNM has been supported by the IMSc Complex Systems Project (12^th^ Plan), and the Center of Excellence in Complex Systems and Data Science, both funded by the Department of Atomic Energy, Government of India. The simulations and computations required for this work were supported by High Performance Computing facility (Nandadevi and Satpura) of The Institute of Mathematical Sciences, which is partially funded by DST.

## SUPPLEMENTARY INFORMATION

### MODELING THE DYNAMICS OF CLOCK GENE EXPRESSION IN CELLS COUPLED VIA NOTCH-DELTA SIGNALING

In the model presented in the main text, we consider a two-component genetic oscillator [58], comprising an activator gene *x* and an inhibitor gene *y*. The parameter values of the model (see Table S1) have been chosen such that it exhibits autonomous oscillatory activity. The interactions between these two genes are schematically shown in Fig. S1 [left]. Each uncoupled oscillator consists of two variables *X* and *Y* that represent the concentrations of the products expressed by genes *x* and *y*, respectively. The trajectory of an uncoupled oscillator in the *X-Y* phase plane is shown in Fig. S1 [right], along with the nullclines 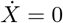 and 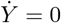. Fig. S1 [left] also displays the nature of the interactions between genes *X* and *y*, where the variables *N^b^*, *N* and *D* describe the Notch-Delta coupling between cells [37].

In a system of coupled oscillators, we observe a variety of synchronization behavior (examples of which are shown in Fig. S2). These can be categorized into 6 principal collective dynamical patterns, namely Inhomogeneous Steady State (ISS), Exact Synchronization (ES), Homogeneous Steady State (HSS), Inhomogeneous In-phase Synchronization (IIS), Anti-Phase Synchronization (APS) and Other synchronization patterns (OTH). In order to classify all observed dynamical patterns into these 6 classes, we use a set of order parameters, as illustrated in the flow chart shown in Fig. S3. We have verified that the results are robust with respect to small changes in the thresholds used to determine these patterns.

We would like to point out that, while in the main text we have exclusively used the variable *Y* to illustrate the dynamical transitions, qualitatively identical behavior can be seen using other variables, such as *N^b^* (as can be seen on comparing Fig. S4 with Fig. 4 of the main text).

**TABLE S1.** Model parameter values used for all simulations in the main text (unless specified otherwise).

| parameter | $a$ | $b$ | $c$ | $e$ | $\beta^N$ | $\gamma$ | $k$ | $\mu$ |
| --- | --- | --- | --- | --- | --- | --- | --- | --- |
| value | 16.0 | 200.0 | 20.0 | 10.0 | 5.0 | 1.0 | 1.0 | 1.0 |

**FIG. S1.**
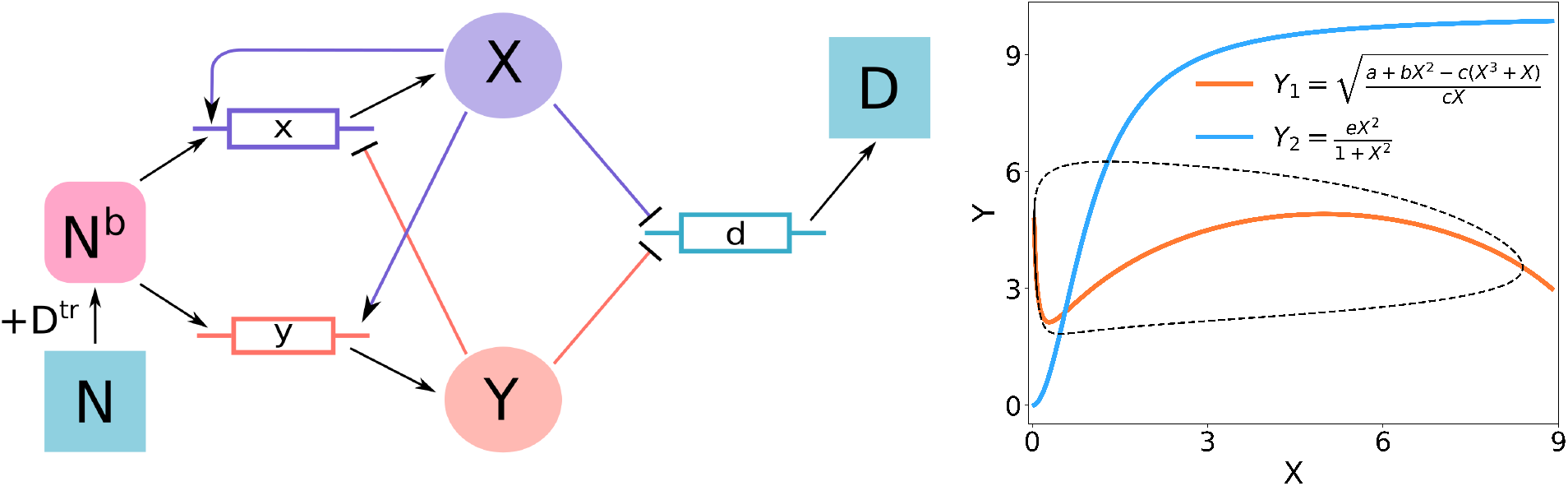
[left] Schematic diagram of the interactions resulting in gene expression oscillations in the model used in the main text. The system comprises an activator gene *x* and an inhibitor gene *y* that yield the gene products *X* and *Y*, respectively. These in turn inhibit the expression of Delta gene *d*, resulting in suppression of the production of Delta ligands *D*. Each cell contains Notch receptors *N* that, when bound with Delta ligands from another cell, causes the Notch Intracellular Domain (NICD, represented through the proxy variable *N^b^*) to cleave off and act as transcriptional factors that upregulate the expression of the downstream genes *x* and *y*. [right] The dynamics of expression levels *X* and *Y* for the activator and inhibitor genes, respectively, in a single uncoupled oscillator, shown in terms of the trajectory of the limit cycle (broken curve) and the nullclines (red: 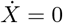, blue: 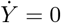) obtained from Eqs. (1)-(2) in the main text, with the parameter values shown in Table S1.

**FIG. S2.**
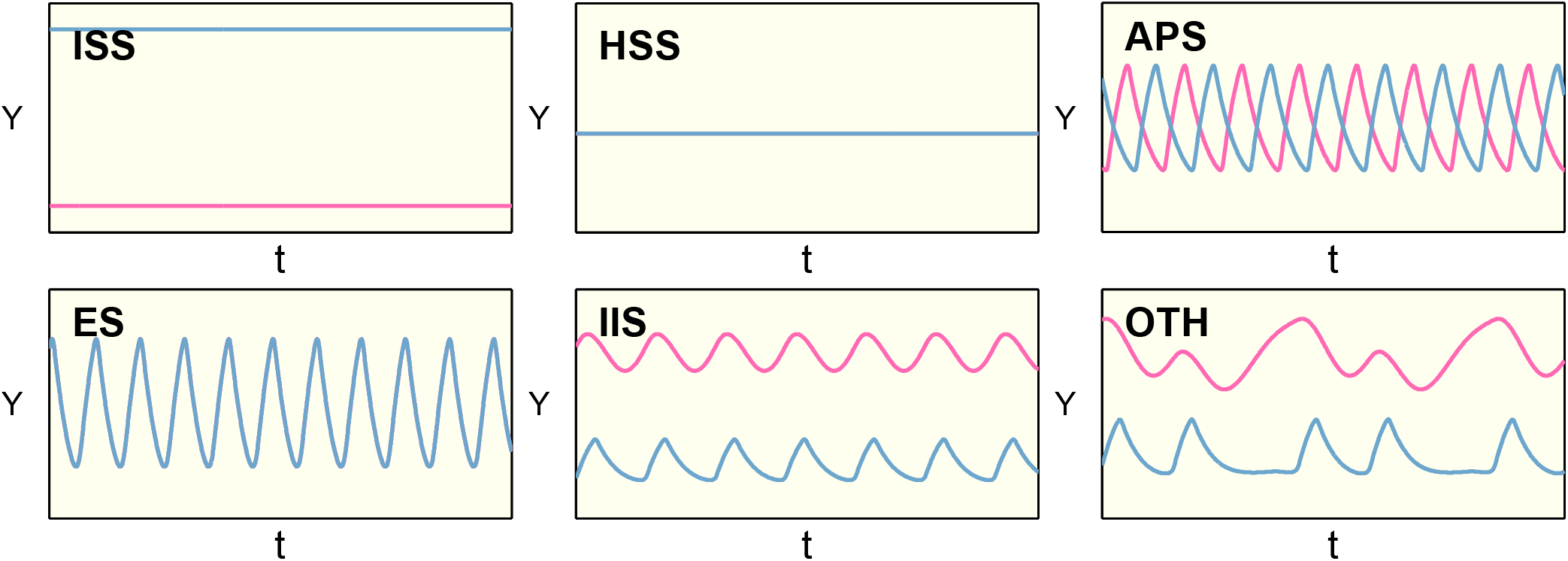
Typical time series of the inhibitor gene expression *Y* for a pair of cells coupled to each other (each exhibiting oscillations in isolation, as shown in Fig. S1). Different dynamical regimes obtained by varying the coupling parameters *f*, *g* and *h* that are shown here correspond to (top row, left) Inhomogeneous Steady State (ISS), (bottom row, left) Exact Synchronization (ES), (top row, center) Homogeneous Steady State (HSS), (bottom row, center) Inhomogeneous In-phase Synchronization (IIS), (top row, right) Anti-Phase Synchronization (APS) and (bottom row, right) Other synchronization patterns (OTH). The activity of the two cells are represented using two different colors in each panel. Note that the two curves overlap in the ES and HSS regimes.

**FIG. S3.**
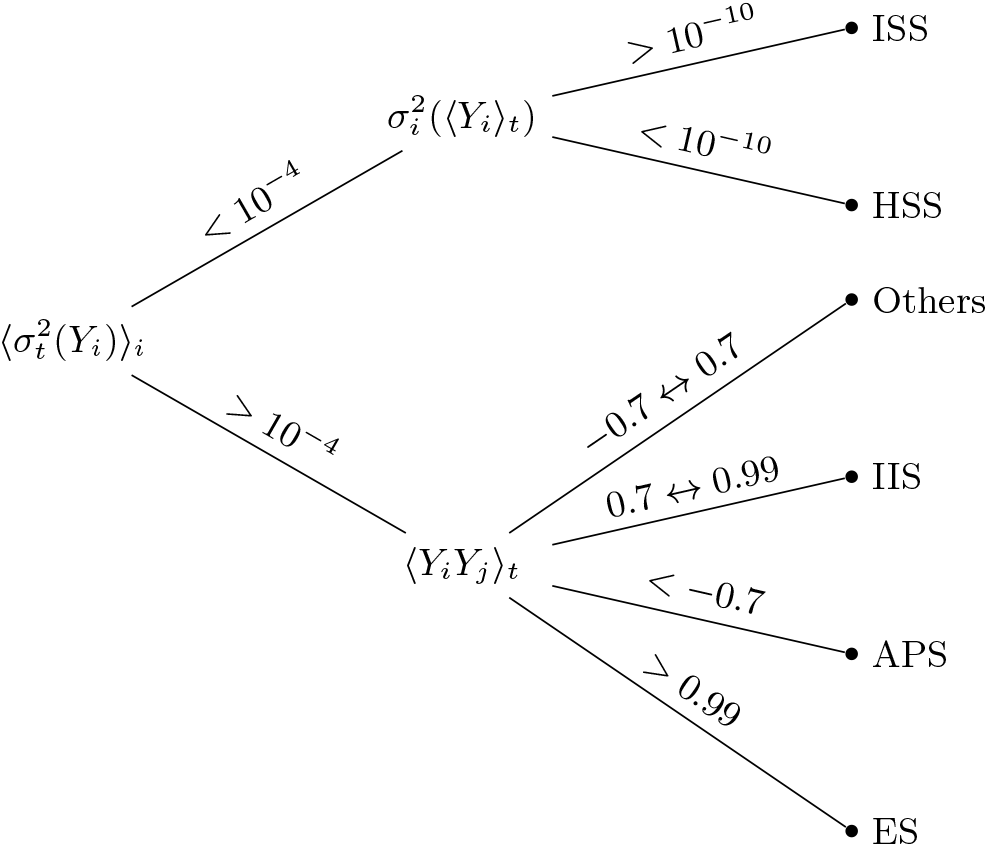
Flow chart illustrating the algorithm used to classify the collective dynamical patterns obtained for a system of two coupled oscillators. To distinguish between oscillating and non-oscillating patterns, we use 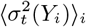 which is the mean temporal variance of the time series of *Y*, calculated over the two oscillators. To determine whether the steady states that the oscillators have reached are the same (corresponding to HSS) or different (corresponding to ISS), we compute the variance of the mean values for the two time series, 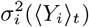. To distinguish between the oscillating patterns, we use the equal time linear correlation between the two time series, 〈*Y_i_Y_j_*〉_*t*_. The classification is robust with respect to small changes in the values of the thresholds, which are displayed in the figure.

**FIG. S4.**
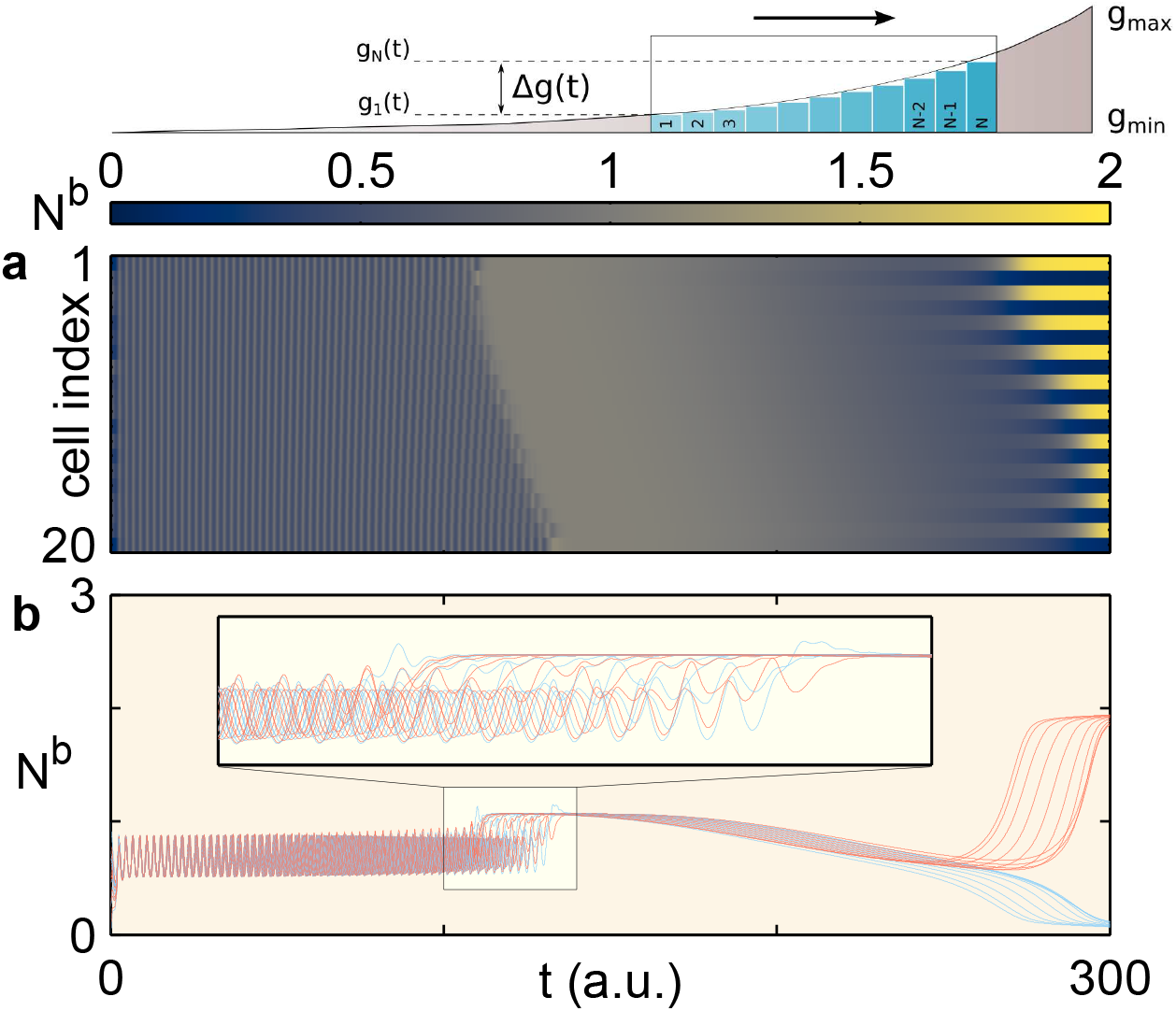
Collective dynamics of a cellular array responding to an exponential gradient of morphogen concentration reproduces the spatio-temporal evolution of PSM activity seen during somitogenesis, similar to Fig. 4 in the main text, but showing bound Notch (*N^b^*) instead of the inhibitor variable *Y*.

### MODELING THE MORPHOGEN GRADIENT IN A GROWING PSM

A crucial element of our model is the gradient of morphogen concentration across the presomitic mesoderm (PSM). We have explicitly considered the morphogen to be Retinoic acid (RA) whose concentration is highest at the anterior end of the PSM. Experimental evidence suggests that as the anteriorly located newly formed somites mature, each of them serve as a source of RA [70–72]. This suggests that a source, diffusion and linear degradation (SDD) model for RA will provide an appropriate description for the variation of the morphogen concentration across space and time. In particular, the source grows in strength as the number of mature somites increases over time. If one considers the PSM to be growing in discrete steps, with each new mature somite (which secretes RA at a rate *ϕ*) being added after a time interval *τ_d_*, the concentration *C_i_* of the morphogen at each putative somite in position *i* on the anteroposterior axis of the growing PSM can be described by the differential equation:

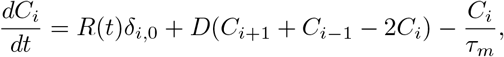

where the time-varying production term *R*(*t*) increases in a step-like manner by *ϕ* after each interval of duration *τ_d_*, *δ*_*i*,0_ is a Kronecker delta function indicating that the source is at the anterior end of the domain (using the simplifying assumption of a point source rather than a distributed one), and *D* and *τ_m_* are the effective diffusion rate and average lifetime of the RA molecules, respectively. We note that qualitatively identical results are obtained if, instead of increasing in discrete time steps, the production rate *R*(*t*) changes in a continuous fashion, for example, as described by the differential equation

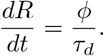

Fig. S5 shows the resulting exponential profile of the morphogen concentration that will be experienced by a particular cell in the expanding PSM, which validates the use of an exponential profile for the morphogen in the main text.

**FIG. S5.**
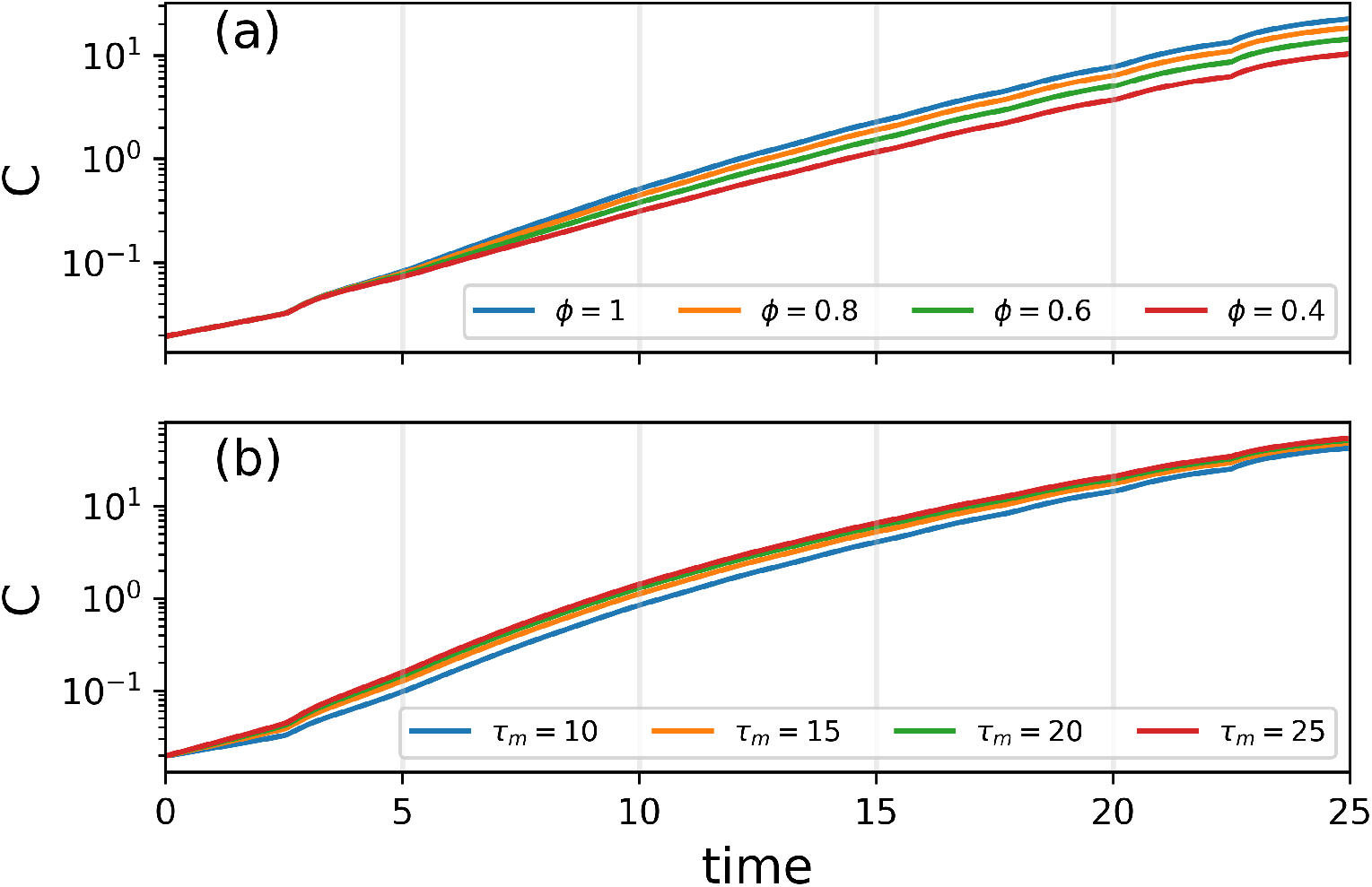
Concentration of the morphogen gradient as sensed by a particular cell in the growing PSM with time, shown for (a) different values of *ϕ* for *τ_m_* = 10, and (b) different values of *τ_m_* for *ϕ* = 0.4. The simulation domain comprises *N* = 10 putative somites at any given time, while the other parameter values are *D* = 1, *τ_d_* = 5 and R(t = 0) = 1.0.

### THE EFFECT OF DELAY ON THE DYANMICS OF COUPLED OSCILLATORS IN THE PRESENCE OF A MORPHOGEN GRADIENT

In the main text, we have described the dynamics of clock gene expressions in cells interacting via Notch-Delta signaling (modulated by a morphogen gradient) where communication delay between the cells is assumed to be insignificant. In Fig. 3 of the main text, we show the dynamical transitions in a pair of coupled cells by focusing on the inhibitor gene product concentration *Y*. We would like to point out that similar dynamics is observed in the other system variables (see Fig. S6). The only exception is the free Notch receptor concentration *N* which, unlike the other variables, grows in an unregulated manner.

In order to investigate the effect of an explicit delay in the system, we incorporate a delay of duration *τ_d_* in both (i) the regulation of *X* and *Y* by *N^b^* and (ii) the repression of the ligand Delta by *X* and *Y*. In Fig. S7, we show the effect of increasing delay, which is expressed in terms of the relative magnitude *τ_d_*/*τ_p_*, where *τ_p_* is the period of the uncoupled oscillator, on the dynamics of the system. We observe the same qualitative nature for the dynamical transitions as seen in Fig. 3 (d) of the main text, suggesting that our results are robust with respect to incorporation of signaling delays.

**FIG. S6.**
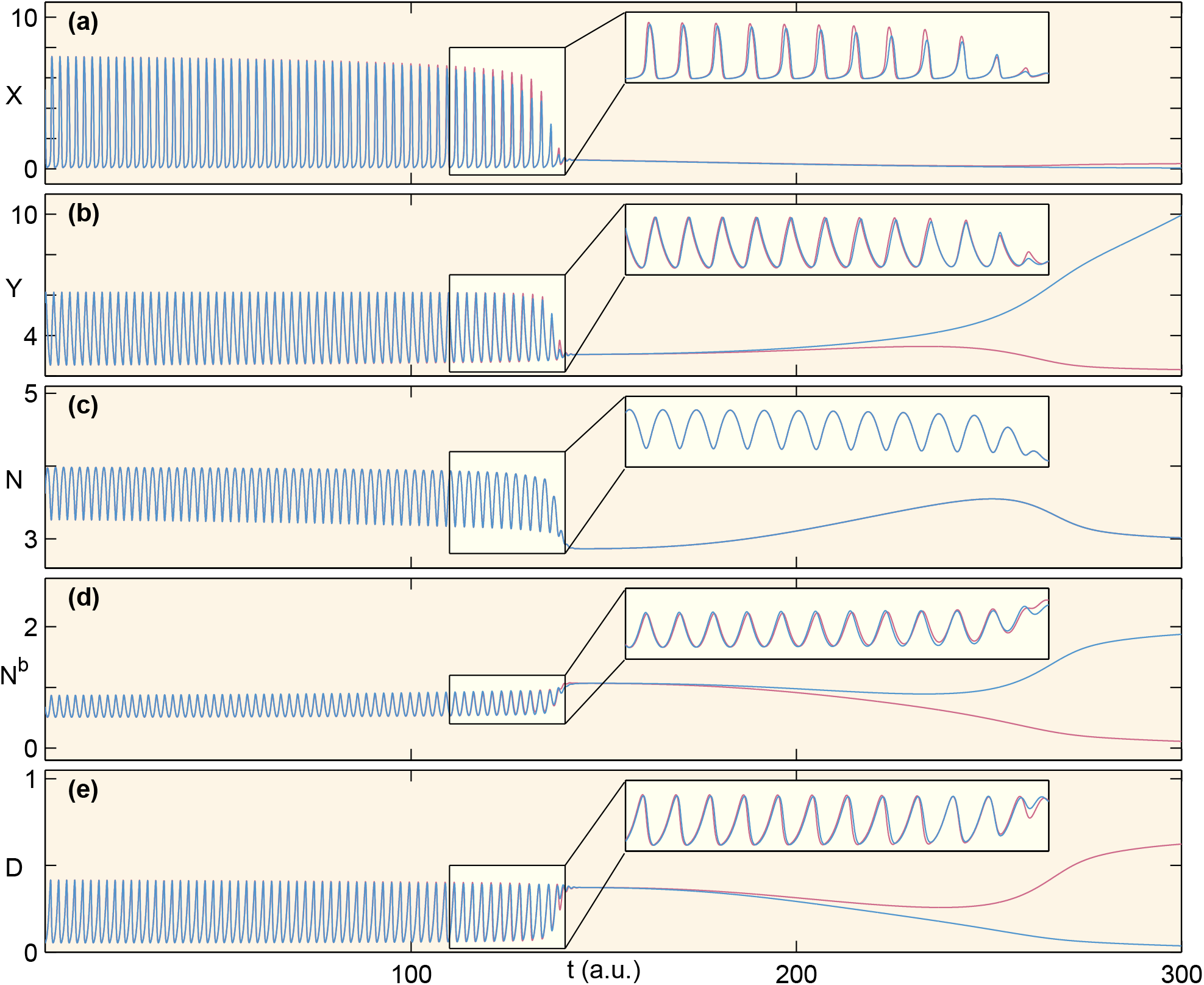
Time series of each of the variables of our model for a pair of coupled oscillators responding to an exponential gradient of morphogen concentration.

**FIG. S7.**
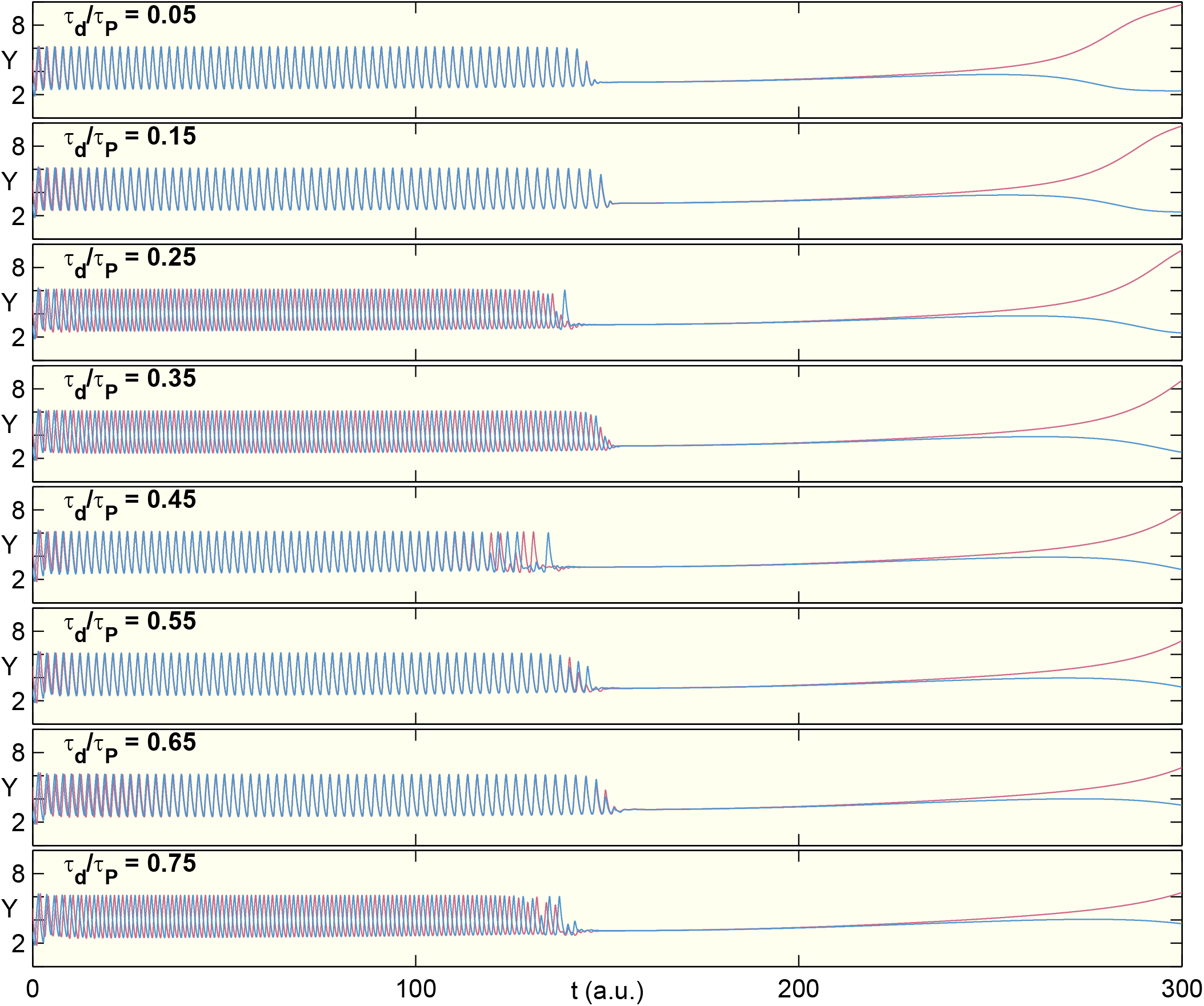
Effect of delay in Notch-Delta mediated interaction on the collective dynamics of a pair of coupled oscillators responding to an exponential gradient of morphogen concentration cell signaling. The effect of the NICD (represented by the proxy variable *N_b_*) on *X* and *Y*, as well as the effect of *X* and *Y* on the expression of the ligand Delta *D*, are each delayed by a period *τ_d_*. The results shown here are obtained for a range of values of *τ_d_*, expressed relative to *τ_p_*, the period of an uncoupled oscillator. We observe that the transition from oscillations to an inhomogeneous steady state is unaffected by the delay.

